# Age-related changes in human skeletal muscle transcriptome and proteome are more affected by chronic inflammation and physical inactivity than primary aging

**DOI:** 10.1101/2023.12.01.569524

**Authors:** Nadia S. Kurochkina, Mira A. Orlova, Maksim A. Vigovskiy, Viktor G. Zgoda, Tatiana F. Vepkhvadze, Nikita E. Vavilov, Pavel A. Makhnovskii, Olga A. Grigorieva, Yakov R. Boroday, Vladislav V. Philippov, Egor M. Lednev, Anastasia Yu. Efimenko, Daniil V. Popov

## Abstract

Evaluation of the influence of primary and secondary aging on the manifestation of molecular and cellular hallmarks of aging is a challenging and currently unresolved issue. Our study represents the first demonstration of the distinct role of primary aging and chronic inflammation/physical inactivity – the most important drivers of secondary aging, in the regulation of transcriptomic and proteomic profiles in human skeletal muscle. To achieve this purpose, young healthy people (n=15), young (n=8) and older (n=37) patients with knee/hip osteoarthritis, a model to study the effect of long-term inactivity and chronic inflammation on the vastus lateralis muscle, were included in the study. It was revealed that widespread and substantial age-related changes in gene expression in older patients relative to young healthy people (∼4,000 genes regulating mitochondrial function, proteostasis, cell membrane, secretory and immune response) were related to the long-term physical inactivity and chronic inflammation rather than primary aging. Primary aging contributed mainly to the regulation of genes (∼200) encoding nuclear proteins (regulators of DNA repair, RNA processing, and transcription), mitochondrial proteins (genes encoding respiratory enzymes, mitochondrial complex assembly factors, regulators of cristae formation and mitochondrial reactive oxygen species production), as well as regulators of proteostasis. It was found that proteins associated with aging were regulated mainly at the post-transcriptional level. The set of putative primary aging genes and their potential transcriptional regulators can be used as a resource for further targeted studies investigating the role of individual genes and related transcription factors in the emergence of a senescent cell phenotype.

## Introduction

Aging causes complex changes in the organism associated with a progressive decrease in functionality at the systemic, organ, tissue, and cellular levels along with the development of various age-related pathologies and an increase in the likelihood of death. Age-related decline in the functionality and mass of skeletal muscles – one of the key integral hallmarks of aging, known as sarcopenia – is related to a decrease in physical performance (endurance) and muscle strength, the ability of muscle tissue to oxidize fats and carbohydrates, and reduction in insulin sensitivity (Campisi et al., 2019; Furrer & Handschin, 2023; Lopez-Otin, Blasco, Partridge, Serrano, & Kroemer, 2023). The decline of skeletal muscle functions affects other systems and organs, leading to the development of various pathologies (metabolic disorders, cardiovascular and nervous system diseases, etc.) and is one of the crucial factors that reduce the quality of life in advanced age.

About a dozen molecular and cellular hallmarks of aging are described in the literature, such as loss of proteostasis, genomic instability, telomere attrition, epigenetic alterations, disabled macroautophagy, deregulated nutrient sensing, mitochondrial dysfunction, cellular senescence, stem cell exhaustion, altered intercellular communication, chronic inflammation, and dysbiosis (Campisi et al., 2019; Lopez-Otin et al., 2023). The appearance of these hallmarks is associated with primary aging, an innate maturational process that occurs with the passage of time from fertilization to death, and secondary aging, which includes the effects of the environment, pathological processes, and/or diseases (Busse, 1969). Evaluation of the influence of primary and secondary aging on the manifestation of molecular and cellular hallmarks of aging is a challenging and currently unresolved issue.

Since age-related changes, especially in humans, are very complex, exploratory studies using high-throughput methods are of the greatest interest. Several studies attempted to identify molecular changes associated with “healthy” aging in human skeletal muscle considering primary aging with minimal contribution of physical impairment and chronic diseases. For this, different muscle molecular parameters were compared between physically active and sedentary elderly people (Hinkley et al., 2023; Lanza et al., 2008) or between young (sedentary) and older people with a comparable level of physical activity/performance (physically active) (Lagerwaard, Nieuwenhuizen, Bunschoten, de Boer, & Keijer, 2021; Lanza et al., 2008; Murgia et al., 2017; Ubaida-Mohien et al., 2019). The last approach compared people of different ages who appeared to had different levels of physical activity throughout their lives; that is, the influence of two factors was simultaneously examined: age and increased levels of physical activity throughout life. The lack of reliable information on the differences between the level of physical activity and performance (and other marks, such as inflammation) in older physically active people compared to the relevant same people at a younger age seems to be a critical limitation of the approach.

The present study aimed to evaluate the role of primary aging and pathological processes/environmental factors including chronic inflammation and physical inactivity in age-related changes of the transcriptomic and proteomic profiles in human skeletal muscle. To achieve this purpose, three groups of volunteers were included in the study: young healthy people (YH), and young and older patients with knee/hip osteoarthritis (YP and OP, respectively). Advanced-stage, symptomatic knee/hip osteoarthritis has been used as a model to study the effect of long-term inactivity/disuse and chronic inflammation on human skeletal muscle (Callahan et al., 2014; Callahan et al., 2015; Miller et al., 2017; Suetta et al., 2007). A comparison of YP *vs*. YH focused on the influence of both chronic inflammation and physical inactivity, while a comparison of OP *vs.* YH (age-related changes in older patients relative to young healthy people) made it possible to evaluate the overall effect of primary and secondary aging on gene and protein expression patterns in skeletal muscle. The comparison of OP and YP (age-related changes in patients occurring against the background of chronic inflammation and physical inactivity) was more relevant to the contribution of primary aging to the observed changes. However, this effect may be markedly modulated by long-lasting (years) chronic inflammation and inactivity, therefore, to identify genes associated with primary aging, the mRNAs and proteins that unidirectionally changed expression were determined when comparing OP *vs.* YP and OP *vs.* YH. Finally, using the transcription factor binding site enrichment analysis, transcriptional factors associated with primary aging genes in human skeletal muscle were predicted.

## Results

### Advanced age- and disease-related physiological changes in subjects

To evaluate the contribution of primary and secondary aging to the molecular changes of human skeletal muscle, three groups of muscle biopsy samples as mentioned earlier were compared (Figure 1A). In older patients (OP) compared to healthy volunteers (YH), there was a pronounced decrease in physical status and an increase in the level of blood leukocytes – a marker of chronic inflammation as the acute inflammatory response was excluded in all subjects before the biopsy (Figure 1B, Supplementary Figure S1). Osteoarthritis-related chronic inflammation and inactivity, as well as age-related sarcopenia, resulted in a two-fold decrease in thigh muscle size, violation in the muscle fiber structure, and a decrease in the total RNA content (Figure 1C and D). In addition, some hallmarks of age-related metabolic disorders were observed in OP: an increase in baseline blood insulin, body mass index, and subcutaneous thigh fat volume (Figure 1D, Supplementary Figure S1).

**Figure 1.**
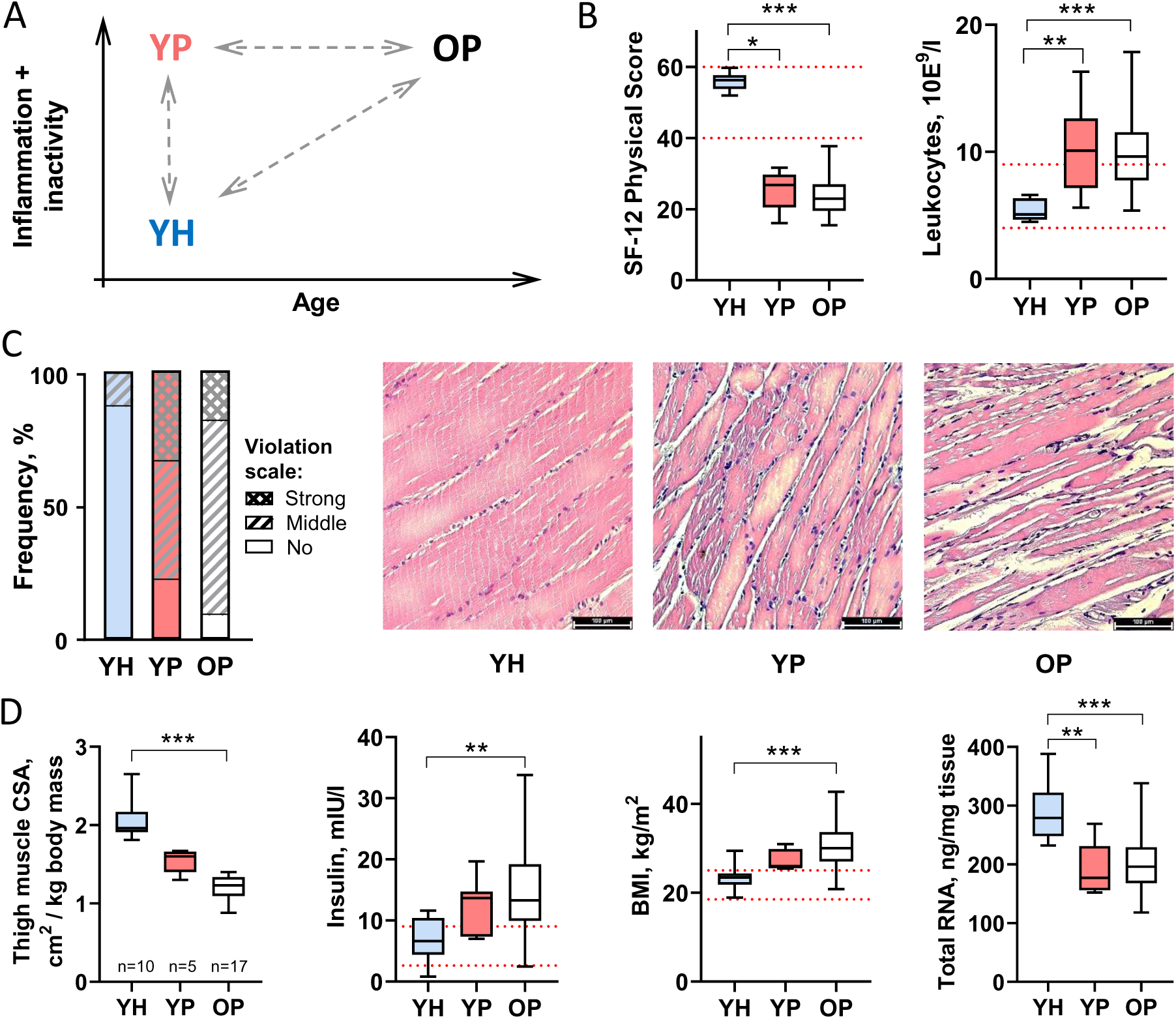
**Pathology- and age-related physiological changes in young (YP) and older (OP) patients with advanced-stage, symptomatic knee osteoarthritis compared to young healthy (YH) individuals.** A – Design of the study. B – Both patient groups demonstrate a substantial decrease in physical status and an increase in the blood leukocyte level, a marker of inflammation. C and D – In patients, chronic physical inactivity and inflammation led to a comparable violation in the muscle fiber structure (C), a progressive decrease in the specific cross-sectional area of thigh muscles (CSA), an increase in the baseline insulin level and body mass index [BMI] (markers of metabolic disorders), as well as to a comparable decrease in the total RNA content in *m. vastus lateralis* (see also Supplementary Figure S1). The red dotted lines indicate the normal range. YH: n = 15, YP: n = 8, OP: n = 37. *, **, and *** – p <0.05, <0.01, and <0.001, respectively.

It is important that young patients (YP) relative to the healthy young control (YP *vs*. YH) showed an increase in the level of blood leukocytes, a decrease in physical status and the content of total RNA in the muscle samples, as well as violation in the muscle fiber structure. However, no significant changes were found in the thigh muscle size, and markers of metabolic disorders in YP, which, apparently, is associated with a shorter duration of the disease (Figure 1B-D, Supplementary Figure S1).

### Physical inactivity and chronic inflammation are key drivers of age-related changes in the skeletal muscle transcriptome in older patients relative to young healthy people

After removing low-expressed genes, >10,000 mRNAs were identified (Supplementary Table S1). Principal component analysis showed that the transcriptomic profile of both patient groups (YP and OP) differed markedly from YH, while the patient profiles were quite similar (Figure 2A). In older patients, widespread changes in the transcriptome profile were revealed relative to young healthy volunteers (OP *vs*. YH): ∼2,000 and ∼2,300 mRNAs were up- and down-regulated, respectively (Figure 2B, Supplementary Table S1). The functional enrichment analysis showed that the changes were strongly associated with a decrease in the expression of genes encoding mitochondrial proteins (>500 mRNAs out of ∼1,000 mRNAs of mitochondrial proteins) and especially respiratory complex proteins, as well as regulators of proteostasis, carbohydrate and fat metabolism, transport and membrane proteins (Figure 2C). Up-regulated mRNAs were associated mainly with secreted, extracellular matrix proteins, and membrane proteins, as well as with immune and inflammatory responses. These findings are in good agreement with previous studies comparing muscle transcriptome in young people with that in older people (Borsch et al., 2021; Su et al., 2015) and in older physically active people, with the level of physical activity comparable to the young control (a model of “healthy” aging) (Lagerwaard et al., 2021).

**Figure 2.**
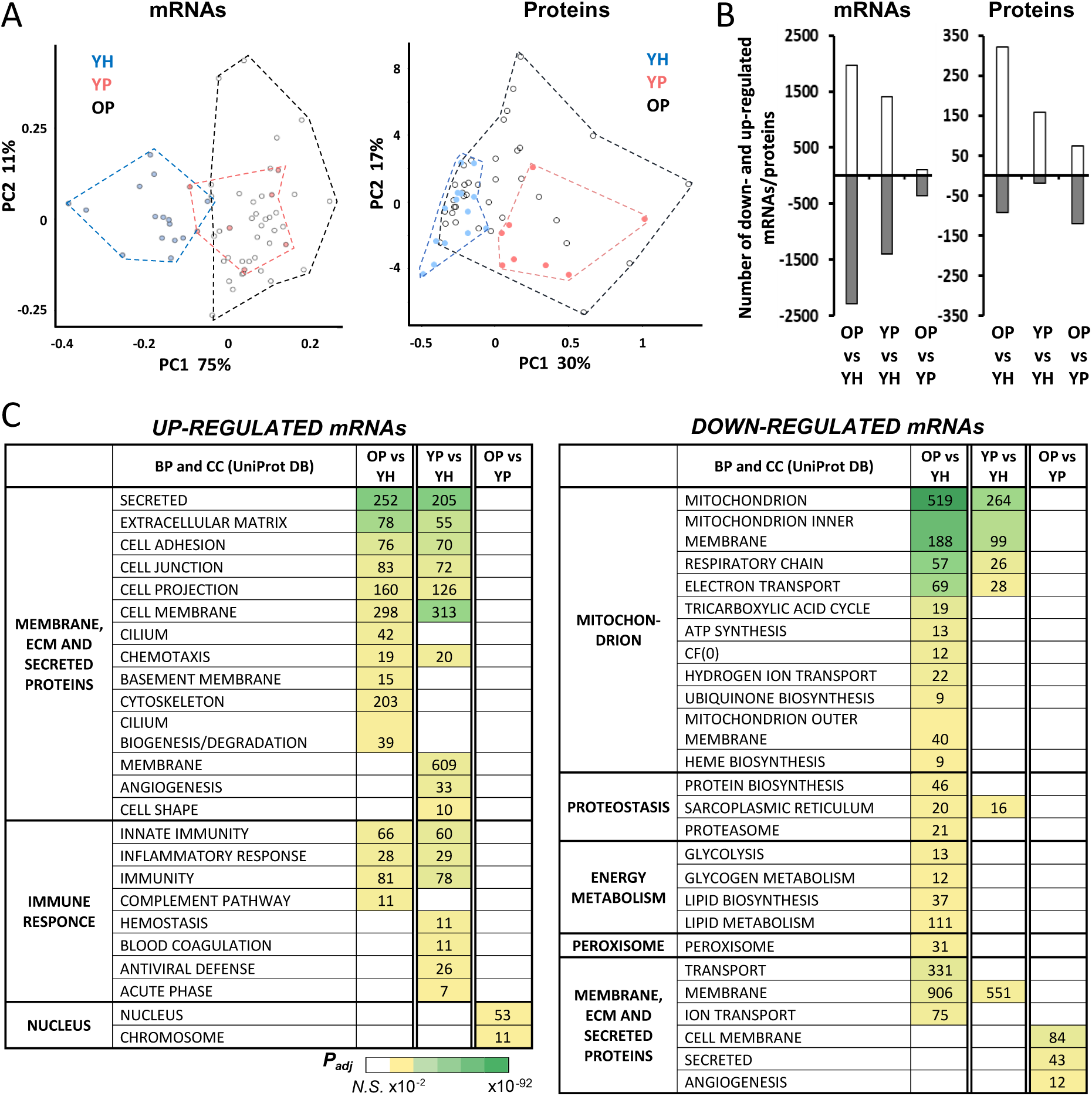
**Chronic inflammation and physical inactivity are key drivers of age-related changes in the skeletal muscle transcriptome in older patients relative to young healthy people.** A – Principal component analysis showed that the transcriptomic and proteomic profiles in young healthy people (YH) were markedly different compared to young patients (YP), while the proteomic profiles in young (YP) and older patients (OP) were much more similar. B – Number of differentially expressed mRNAs and proteins for each comparison. C – Function enrichment analysis revealed that chronic inflammation and physical inactivity were key drivers of age-related changes in the skeletal muscle transcriptome. BP and CC, biological processes and cellular components, respectively; the heat map shows the significance level (NS – non significant; see also Supplementary Table S2). YH: n = 15, YP: n = 8, OP: n = 37.

The number of differentially expressed genes in YP relative to YH was 1.5 times less than in OP *vs*. YH (∼1,400 and ∼1,400 up- and down-regulated mRNAs; Figure 2B, Supplementary Table S1), however, the direction of the transcriptome profile changes was similar (Supplementary Figure S2, Figure 2C). In particular, there was a decrease in the expression of many genes encoding mitochondrial and membrane proteins, as well as sarcoplasmic reticulum proteins, and an increase in gene expression of secreted, membrane, and extracellular matrix proteins, and regulators of inflammatory and immune response (Figure 2C). Curiously, in OP the number (Figure 2C) and expression (Supplementary Figure S3) of up-regulated mRNAs enriched functional categories “inflammatory response” and “immunity” were lower than in YP, which may indicate age-related suppression of the proinflammatory response in the patient’s muscle.

Transcriptomic changes in older patients relative to younger patients (OP *vs*. YP) were more likely associated with primary aging than those between other groups; however, it is important to note that these changes occurred against the background of long-lasting (years) inflammation, which could significantly modulate the transcriptome profile (Supplementary Figure S3). Surprisingly, these transcriptomic changes (OP *vs*. YP) were modest (∼110 and ∼360 up- and down-regulated mRNAs, respectively; Figure 2B, Supplementary Table S1) and fundamentally different from the previous comparisons: the up-regulated mRNAs enriched the functional categories “nucleus” and “chromosome”, while down-regulated mRNAs enriched the terms “cell membrane” and “secreted” (Figure 2C, Supplementary Figure S2, Supplementary Table S2). Altogether, the obtained transcriptomic data, as well as a pronounced decrease in the total RNA content in YP relative to YH, indicate the key role of chronic inflammation and inactivity in the development of age-related changes in the skeletal muscle transcriptome in OP relative to YH.

### Proteins with distinguished functions are differently regulated at the mRNA level, while aging-related protein changes in patients are regulated mainly at the post-transcriptional level

Sarcomeric proteins make up about half of muscle proteins, which limits the depth of shotgun proteomic analysis in this tissue. Proteins highly abundant within skeletal muscle were mainly detected (n=1,899; sarcomeric and mitochondrial proteins, energy metabolism enzymes, chaperones, etc.), of which 1,387 proteins were detected in all samples (n=60) (Supplementary Table S3). Principal component analysis showed that the YH proteomic profiles, like the transcriptomic profiles, markedly differed compared to YP, while the YP and OP proteomic profiles were substantially more similar (Figure 2A). As expected, the greatest proteomic change (413 proteins) was found in older patients relative to young healthy individuals (OP *vs*. YH; Figure 2B). In contrast to the transcriptomic data, the number of differentially expressed proteins in OP *vs*. YP (196 proteins) was comparable to YP *vs*. YH (178 proteins; Figure 2B), which may be due to the fact that only a fraction (n=1,899, less than a third) of proteins expressed in human tissues (∼10,000 (D. Wang et al., 2019)) was detected.

It was determined that only a third of age- and pathology-related changes in protein content (OP *vs*. YH and YP *vs*. YH, respectively) correlated with changes in their mRNA level (sets *“protein UP-mRNA UP”* and *“protein DOWN-mRNA DOWN”* in Figure 3A). Moreover, when comparing older patients to young patients (OP *vs*. YP), changes in the content of almost all proteins occurred without changes in the expression of the corresponding mRNAs (sets *“protein UP-mRNA NS”* and *“protein DOWN-mRNA NS”* in Figure 3A), meaning that age-related protein changes in patients were regulated mainly at the post-transcriptional level.

**Figure 3.**
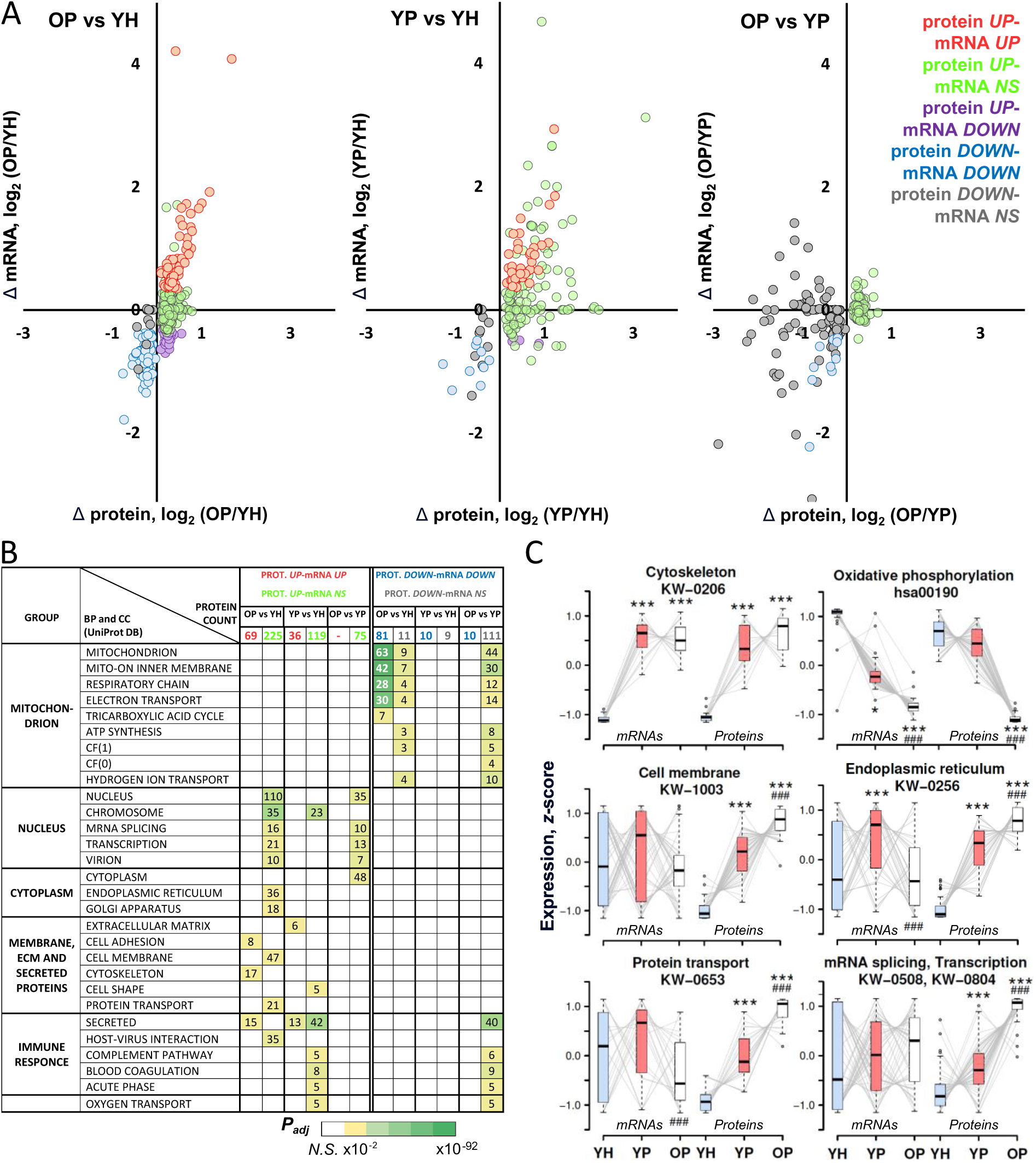
**Proteins with different functions are differently regulated at the mRNA level, while aging-related protein changes in patients are regulated mainly at the post-transcriptional level.** A – Changes in the content of highly abundant proteins are weakly regulated at the mRNA level, especially in the patient groups (OP *vs.* YP). Different colors represent the protein regulation pattern at the mRNA level. NS – non significant. B – Function enrichment analysis revealed specific proteomic changes for each comparison. BP and CC, biological processes and cellular components, respectively; the heat map shows the significance level (NS – non significant; see also Supplementary Table S4). C – Analysis of the intergroup expression trajectory (YH-YP-OP) of differentially expressed proteins belonging to some enriched functional categories and their mRNAs showed that pathology- and age-related changes in the abundance of protein with various functions were differently regulated at the mRNA expression level. Normalized expression (*z*-score) for each protein/mRNA, as well as the name and ID of functional categories, are indicated. * – difference from YH, # – difference from YP: one and three symbols – p <0.05 and <0.001, respectively. YH: n = 15, YP: n = 8, OP: n = 37.

A comparison of proteomic and transcriptomic data (Figure 2C, 3B and C, Supplementary Table S4) showed that age-related changes in the content of proteins with various functions were differently regulated at the level of their mRNA expression, namely, intergroup expression trajectory (YH-YP-OP) of proteins and mRNAs were significantly different. In particular, the expression of cytoskeletal proteins correlated with the expression of their mRNAs in all comparisons. On the contrary, a decrease in the expression of genes encoding oxidative phosphorylation enzymes – the most pronounced change in the transcriptomic profile – did not cause a decrease in the content of corresponding proteins in young patients (YP *vs.* YH), but correlated with a decrease in these proteins in older patients (OP *vs.* YP and OP *vs.* YH). Other proteins (proteins of the cell membrane, endoplasmic reticulum, regulators of protein transport, splicing, and transcription) showed a pathology- and age-related increase in their content, which occurred against the background of multidirectional regulation of their mRNAs (Figure 3C).

### Genes associated with primary aging in human skeletal muscle

It was found that in older patients, relative to young patients (OP vs. YP), the direction of changes in the expression of many genes did not coincide with those observed in older patients relative to the healthy control (OP *vs.* YH; Figure 3C, Supplementary Figure S3). This may partly be explained by the fact that in younger patients, up-regulated genes are more enriched immune/inflammatory functional categories than in older patients (at the mRNA and protein levels; Figure 2C and 3B, respectively), which indicates muscle inflammation. Therefore, genes associated with primary aging were defined as genes whose proteins (or mRNAs, since it was possible to determine the expression of less than a third of all muscle proteins) unidirectionally changed expression in OP *vs.* YP and OP *vs.* YH (mRNAs whose protein products did not change expression were excluded) (Figure 4A, Supplementary Figure S2), and then their functions and potential protein-protein interactions were described (Figure 4B, Supplementary Table S5). The largest clusters of interacting proteins were formed by mitochondrial and nuclear proteins. Almost all genes of mitochondrial protein decreased expression; these included enzyme subunits of all respiratory complexes, mitochondrial complex assembly factors (such as *TIMMDC1* and *SDHAF3*, and *COA7*), regulators of cristae formation (*IMMT*, *SAMM50*, *APOO*, and *CHCHD3*), phosphate, ADP, and ATP carriers (*CKMT2*, *SLC25A3* and *-4* [or *ADP/ATP translocase 1*]), subunits of the pyruvate dehydrogenase complex (*PDHA1* and *PDHB*), and citrate cycle enzymes (*SUCLA2* and *SUCLG1*). On the contrary, the genes of the largest cluster of nuclear proteins – DNA damage and repair genes (*DDB2*, *RADX*, *MRNIP*, etc.), as well as RNA processing regulators (*SRSF2*, *HNRNPA1*, *-A2B1*, *-C*, *-M*, *-K*, etc.) – were predominantly activated. In addition, there are a large number of up- and down-regulated transcription regulators including transcription factors and coactivators playing a role in mitochondrial biogenesis (*PPARGC1A* [*PGC1a*]) and muscle differentiation (*EID2B*), as well as genes encoding ubiquitin system proteins (*UBAP1L*, *COP1*, *WWP1*, etc.). Other large groups included: predominantly up-regulated folding regulators (*HSPA2* [*HSP70*], *DNAJ5B,-C5B* [*HSP40s*], small chaperones *HSPB2* and *-B3*, etc.), as well as multi-directionally regulated various transporters (*RAB21*, -*2A*, *STAU2*, *SLC14A1*, -*26A11*, etc.), growth factors and immunomodulators (*IGF1*, *HDGF*, *IL17RC*, *VSIR*, etc.), kinases/phosphatases (*RSKR*, *ERBB2*, *PUDP*, *PPM1A*, etc.), translation (*EIF4H*, *ETF1*, etc.) and cytoskeletal (*SYNM*, *ABI2*, and *FAM171A1*) regulators. Then it was revealed that a third of the primary aging genes identified in the study were previously associated with age-related changes in gene expression in muscle and other tissues (Supplementary Figure S4), which partially validated the findings of the present study. Interestingly, we revealed that the set of primary aging genes was significantly enriched in genes that changed expression in skeletal muscle of older individuals/animals (Su et al., 2015), but not in other tissues (Avelar et al., 2020; Palmer, Fabris, Doherty, Freitas, & de Magalhaes, 2021) and senescent human fibroblasts (Casella et al., 2019), an *in vitro* model for studying the effects of primary aging (Supplementary Figure S4). The last finding indirectly indicated that the primary ageing genes identified in our study are specific to post-mitotic multinucleated muscle fibers.

**Figure 4.**
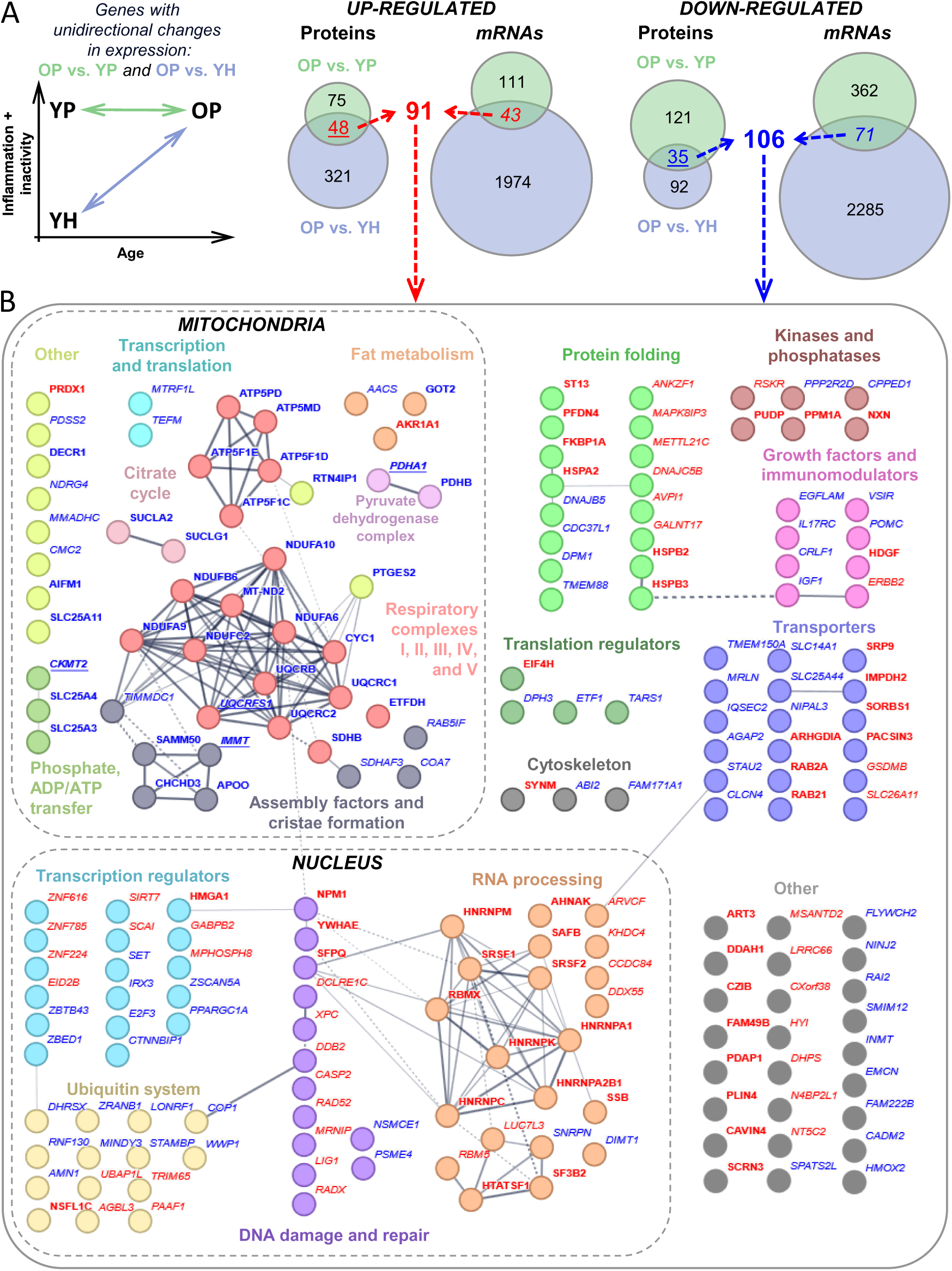
**Genes associated with primary aging.** A – Genes associated with primary aging were identified as proteins (as well mRNAs) that unidirectionally change expression in both OP *vs.* YP and OP *vs.* YH (YH: n = 15, YP: n = 8, OP: n = 37). B – Functions of primary aging associated genes. Analysis of protein-protein interactions and search of the GO and KEGG BRITE databases revealed the major functional groups of proteins, including proteins interacting with each other. Proteins and mRNAs are shown in *italics* and **bold**, respectively: underlining indicates changes in both mRNA and protein level. Up- and down-regulated proteins or mRNAs are shown in red and blue, respectively (see also Supplementary Table S5). Thicker edges indicate higher protein-protein interaction scores.

### Putative transcription factors regulating primary aging gene expression in human skeletal muscle

A change in gene expression profile can be caused by changes in cell signaling, proteostasis, epigenetic regulation, etc. Identifying the mechanisms (in particular, transcription factors) that control the expression of primary aging genes is of great importance. To predict transcription factors involved in the regulation of primary aging genes, the individual promoter regions (see Methods) of primary aging genes (43 and 71 up- and down-regulated mRNAs in Figure 4A) were examined using the transcription factor binding site enrichment analysis. Significant enrichment was found only for the promoters of down-regulated mRNAs, which indicated a decrease in the activity of the putative transcription factors. On the other hand, down-regulation of these mRNAs may be due to the fact that half of the predicted factors (NR3C1, ETS1, NR2F6, ZNF436, BCL6, GATAD2A, ZNF461) can act as repressors (Figure 5, Supplementary Table S6). The highest enrichment was found for transcription factor motifs: ZFP64, NR3C1, ETS1, NR2F6, ZNF436, ZNF528, and a dimer RELB:NFKB1.

**Figure 5.**
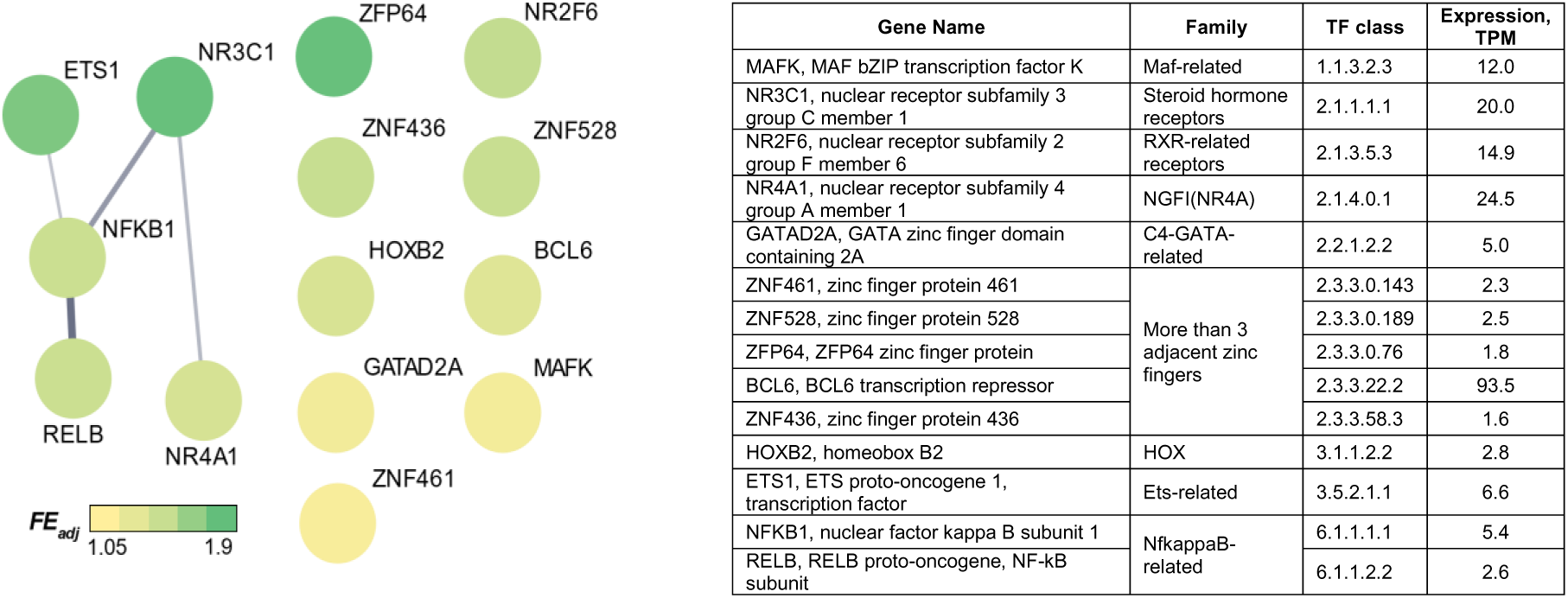
**Transcriptional factors associated with down-regulated primary aging genes (mRNAs).** The heat map shows adjacent fold enrichment (FE_adj_, statistically corrected odds ratios with a confidence interval of 99%) for transcription factor binding site frequency in individual promoters of primary aging genes relative to that of random genes showing no differential expression (see also Supplementary Table S6). Thicker edges indicate higher protein-protein interaction scores.

## Discussion

The present work examined, for the first time, the role of pathological processes (chronic inflammation and physical inactivity) and primary aging in age-related changes of the transcriptomic and proteomic profiles in human skeletal muscle. As expected, widespread changes were found in gene and protein expression profiles in older patients relative to young healthy individuals (associated mainly with the suppression of mitochondrial protein gene expression and increased expression of extracellular matrix, secreted, immune, and nuclear protein genes; Figure 2 and 3, Supplementary Table S2 and S4), which was in good agreement with studies comparing older people (Borsch et al., 2021; Cutler et al., 2017; Lanza et al., 2008; Su et al., 2015), as well as older physically active people (a model of “healthy” aging) (Lagerwaard et al., 2021; Lanza et al., 2008; Murgia et al., 2017; Ubaida-Mohien et al., 2019), with young subjects. Next, it was revealed that young patients showed significant and similar to the previous comparison changes in the phenotype (an increase in markers of inflammation and a violation in the muscle fiber structure), as well in transcriptome relative to the young healthy control (Figure 1 and 2), which clearly indicated that physical inactivity and chronic inflammation could be considered key drivers of age-related changes in skeletal muscle of older patients (relative to young healthy people).

To identify genes associated with primary aging, genes that unidirectionally changed the expression of proteins (or mRNAs) in OP *vs.* YP and in OP *vs.* YH (Supplementary Figure S2, Figure 4) were determined. Unexpectedly, only a few of such genes were revealed compared to pathology-associated genes (YP *vs.* YH). The lack of overlapping of primary aging genes and pathology-associated genes (Supplementary Figure S2) clearly indicates the distinct functions of these gene sets. It is possible that a small number of primary aging-related genes are specific for skeletal muscle only (Supplementary Figure S4В). A number of studies showed that the age-related acquisition of senescence markers varied significantly in cells of different mammalian tissues, while in skeletal muscle this increase [such as DNA damage foci-associated protein H2AX (C. Wang et al., 2009), senescence-associated heterochromatin foci-associated core histone macro-H2A.1 (H2AY) (Kreiling et al., 2011), telomere dysfunction induced foci-associated TP53-binding protein 1 (TP53B) (Jeyapalan, Ferreira, Sedivy, & Herbig, 2007), and tumorigenesis (Seely, 1980)] is lower and/or absent compared to cells of other tissues (hepatocytes/liver, dermis and fibroblasts/skin, alveoli/lung, lymphocytes/spleen, crypt/small intestine). A small number of primary aging genes (as well as a slight age-related increase in senescence markers) might be a specific characteristic of muscle fibers – fully differentiated postmitotic (non-replicated) cells with a very high plasticity potential. The main function of skeletal muscles is contractile activity; maintaining/increasing the level of physical activity in the elderly was shown to have a pronounced positive effect on the content and function of skeletal muscle mitochondria, fat and carbohydrate metabolism, insulin sensitivity, and endurance performance (Distefano et al., 2018; Lanza et al., 2008) and can induce an increase in muscle mass and strength (Papa, Dong, & Hassan, 2017; Peterson, Rhea, Sen, & Gordon, 2010), thereby contributing to the exercise-induced increase in lifespan (Campisi et al., 2019; Goh, Wong, Soh, Maier, & Kennedy, 2023). This suggests that senescent skeletal muscle cells (unlike many other somatic cells) retain an exceptionally high potential to restore their functions under the action of a specific physiological stimulus – regular contractile activity. This reasoning is supported by the fact that some of the molecular “anti-aging” effects of long-term physical training (transcription and methylation reprogramming) coincide with that of partial *in vivo* reprogramming by short-term induction of the Yamanaka transcription factors (Jones et al., 2023), facilitating cellular plasticity and regeneration that are characteristics of a more youthful cell state (Gill et al., 2022; Ocampo et al., 2016). Together, these data indicate the need to identify primary aging genes in cells of various tissues and compare them with skeletal muscle. This approach may reveal mechanisms of sensitivity/resistance to the emergence of a senescent cell phenotype and also indicates the possibility of using skeletal muscle as a model to study restoring the functions of senescent cells.

It was revealed that only a third of age-related changes in the content of muscle proteins (predominantly highly abundant proteins) correlated with changes in their mRNA level (OP *vs.* YH in Figure 3A), which is consistent with the work that investigated age-related changes in proteomic and transcriptomic profiles in mouse skeletal muscle (Hunt et al., 2021). A similar pattern was observed for pathology-related changes (YP *vs.* YH in Figure 3A), meaning that changes in the abundance of the majority (more than half) of the detected proteins are regulated by changes in the rate of transcript-specific translation and/or protein-specific degradation. This conclusion complements studies showing that the age-related decrease in the fractional protein synthesis rate in human skeletal muscle (Breen et al., 2011) occurs together with transcript-specific deregulation in translation efficiency (ribosome occupancy), which decreases for genes encoding mitochondrial proteins in human skeletal muscle (Tharakan, Ubaida-Mohien, Piao, Gorospe, & Ferrucci, 2021), mitochondrial, membrane proteins and translation regulators in mouse liver and kidney, but increases for genes encoding inflammatory proteins (Anisimova et al., 2020). Curiously, the change in the content of almost all proteins in patients (OP *vs.* YP in Figure 3A), as well as proteins associated with primary aging (for example, mitochondrial proteins, Figure 4B, Supplementary Table S5), occurs without changing the expression of their mRNAs, indicating that (primary) aging-related changes in expression of highly abundant proteins are exclusively regulated at the post-transcriptional level (translation and protein degradation). In addition, it was demonstrated that proteins with different functions were differently regulated at the mRNA level and the pattern of this regulation was modulated by aging and the presence of pathological conditions (inactivity/inflammation) (Figure 3). The specificity in the regulation of proteins with different functions was shown previously by high-throughput approaches in mouse fibroblasts (Schwanhausser et al., 2011), human hematopoietic stem cells differentiating into erythroid progenitors (Liu et al., 2017), and human skeletal muscle after a period of physical training (Makhnovskii et al., 2020). Overall, these results clearly demonstrate the presence of conserved function-specific mechanisms that regulate protein abundance under the influence of various stimuli, including chronic inflammation/inactivity and aging.

Analysis of the functions of putative primary aging genes identified in this study (Figure 4B, Supplementary Table S6) showed that ∼40% of them were previously attributed to aging and/or senescence in muscle and other tissues (Supplementary Figure S4). For example, *PPARGC1A,* an important regulator of mitochondrial biogenesis and antioxidant defense (Ji & Kang, 2015; Wenz, Rossi, Rotundo, Spiegelman, & Moraes, 2009), genes encoding mitochondrial proteins (subunits of the pyruvate dehydrogenase complex and all respiratory complexes) ((Murgia et al., 2017; Ubaida-Mohien et al., 2019) (all down-regulated), regulators of splicing (Lee et al., 2016; Ubaida-Mohien et al., 2019), and DNA repair (Schumacher, Pothof, Vijg, & Hoeijmakers, 2021) (predominantly up-regulated). It is important to note that many genes in these groups were not previously associated with aging and are therefore of interest for future research. Interestingly, down-regulated genes encoding assembly factors and regulators of cristae formation, as well as chaperones and associated proteins, were identified as primary aging genes, which is in line with data on age-related disturbances in the cristae structure (Beregi, Regius, Huttl, & Gobl, 1988) and proteostasis (Goh et al., 2023) in old muscles. Other interesting examples: the down-regulation of *IGF1*, a growth factor regulating muscle protein expression/synthesis, is consistent with its decreased level in plasma in people with sarcopenia (Bian et al., 2020) and expression in various tissues of long-lived species (Tyshkovskiy et al., 2023), as well as the down-regulation of mitochondrial creatine kinase (*CKMT2*) and ADP/ATP translocase 1 (*SLC25A4*), those recently identified (along with mitochondrial-bound hexokinase I or II) as key regulators of mitochondrial reactive oxygen species production in skeletal muscle and other tissues (Vyssokikh et al., 2020). Identifying the mechanisms regulating the expression of primary aging genes is of fundamental importance. Transcription factors associated with down-regulated (at the mRNA level) primary aging genes were predicted (Figure 5). The validity of the prediction of some transcriptional regulators (such as NFKB1, BCL6, NR4A1, NR2F6) is confirmed by previous mechanistic studies. Namely, knockout of *Nfkb1* leads to early mice aging, which is associated with reduced apoptosis and increased cellular senescence (Bernal et al., 2014; Jurk et al., 2014). Knockdown of *Bcl6*, a terminal differentiation factor for C2C12 myoblasts, induces apoptosis (Kumagai et al., 1999). Additionally, orphan nuclear receptors of the NR4A family are considered age-dependent regulators of fat and carbohydrate metabolism and DNA repair (Paillasse & de Medina, 2015). On the contrary, overexpression of the repressor *Nr2f6* suppresses the expression of *Ppargc1a*, genes related to energetic metabolism and muscle contraction, induces muscle atrophy, and impairs muscle force production (Guimarães et al., 2023).

## Limitations

The lack of a “healthy” older group (physically active older persons with minimal contribution of physical impairment and chronic diseases) limits the interpretation of our data. Comparison of this group with older patients would be useful to identify genes associated with inflammation and decreased physical activity in older patients. It can be assumed that this set of genes differs from that identified for YP *vs.* YH. Another limitation of the study is the small number of people in the young patient group. Further studies with an increase the number of compared groups and sample size seem to be an interesting and promising direction.

Recent studies found some sex-related differences in human skeletal muscle transcriptome during ageing (de Jong et al., 2023; Huang et al., 2023). We analyzed sex-related differences in the older patient group (32 old females vs. 5 old males) and found small number (43) of differentially expressed genes and little overlap of this genes with other comparisons (OP *vs.* YH and OP *vs.* YP). Moreover, no differentially expressed proteins were found for this comparison (Supplementary Figure S5). Together, this suggests little influence of sex-related differences in gene expression on the main results of our study.

## Conclusions

To the best of the authors’ knowledge, this study represents the first demonstration of the distinct role of primary and secondary aging in the regulation of transcriptomic and proteomic profiles in human skeletal muscle. Using the specific approach to the selection of experimental groups, it was revealed that widespread and substantial age-related changes in gene and protein expression (genes regulating mitochondrial function, proteostasis, cell membrane, secretory and immune response) in older patients relative to young healthy people were related to the long-term physical inactivity and chronic inflammation rather than primary aging. The impact of primary aging was also separated, and it was demonstrated that it contributed mainly to the regulation of genes encoding nuclear proteins (including genes related to DNA repair, RNA processing, and transcription regulation), mitochondrial proteins (genes encoding oxidative metabolism enzymes, mitochondrial complex assembly factors, regulators of cristae formation and mitochondrial reactive oxygen species production), as well as regulators of proteostasis. Comparing the changes in the expression of proteins and the corresponding mRNAs, it was found that proteins associated with aging were regulated mainly at the post-transcriptional level. The study of this regulatory mechanism seems to be a promising avenue for future investigations. The set of putative primary aging genes and their potential transcriptional regulators identified in this study can be used as a resource for further targeted studies investigating the role of individual genes and related transcription factors in the emergence of a senescent cell phenotype. The findings of this study could also provide novel opportunities for the diagnostics and therapy of sarcopenia and other age-associated diseases.

## Methods

The study was conducted in accordance with the Declaration of Helsinki and approved by the Ethics Committee of the Medical Research and Educational Center of Lomonosov Moscow State University (IRB00010587, Protocol No. 2/20 of March 16, 2020). Informed consent was obtained from all subjects involved in the study.

### Study design

The study involved 37 older patients with advanced-stage, primary symptomatic knee/hip osteoarthritis (OP; median age 72 years and interquartile range [69–77] years, M:F = 5:32), 8 young patients with the same diagnosis (YP; 39 [37–42] years, M:F = 7:1), and 15 young healthy volunteers (YH; 35 [28–38] years, M:F = 13:2). Exclusion criteria were: 1) history of cancer or systemic diseases; 2) mental, physical and other reasons that do not allow to adequately assess the patient’s behavior and correctly comply with the conditions of the study protocol; 3) pregnancy and lactation. We calculated the power of the study for RNA-seq data using the PROPER R package and simulation-based method to compute power-sample size relationship (Wu, Wang, & Wu, 2015). It was shown that *i)* the sample size n >7 and the cut-off criteria used in the study (see below) yielded the power >0.8 and *ii)* the number of differentially expressed genes in this case was independent of sample size (Supplementary Figure S6).

Venous blood samples were taken after an overnight fast to determine the content of leukocytes, neutrophils (XN-2000 hematology analyzer, Sysmex, Japan), total cholesterol, glucose, and insulin (Cobas 6000 immunochemical analyzer, Roche, Germany) to estimate HOMA IR, then the physical status of the volunteers (the SF-12 questionnaire that evaluates such parameters as physical functioning, physical role functioning, bodily pain and general health (Ware, Kosinski, & Keller, 1996)) and thigh muscle size (see below) were examined. Biopsies of muscle tissue from *m. vastus lateralis* were taken after an overnight fast for histological examination, transcriptomic and proteomic studies.

### Thigh muscles size

The muscle and subcutaneous fat thigh cross-sectional area of both legs was assessed using computed tomography (Somatom Scope, Siemens, Germany) and the RadiAnt DICOM Viewer (Medixant, Poland) on three cross-sections located in the middle between the knee joint space and the center of the femoral head, as well 1 cm distal and proximal of this cross-section. Then, the average cross-sectional area of the thigh muscles was normalized to body mass.

### Muscle biopsy

In young healthy voluntaries, samples from *m. vastus lateralis* were taken under local anesthesia (2 ml 2% lidocaine) using a modified Bergstrom needle with aspiration; in patients – immediately before knee/hip replacement surgery. All patients were in hospital at least two days before the biopsy study. The healthy participants were asked to refrain from any physical exercise for 24 hours prior to visiting the laboratory and to minimize physical activity on the way to the study center. Biopsies were taken after 10-12 h overnight fast between 9:00 – 12:00. All samples were placed in the Custodiol buffer (Dr. Franz Köhler Chemie GmbH, Germany) and processed within 10–15 minutes after the biopsy. Thus, visible fragments of connective and adipose tissue were removed; a part of the tissue was fixed for histological examination, and another part was frozen in liquid nitrogen and stored at-80 °C.

### Histological study

Samples of muscle tissue were fixed with 10% neutral formalin and then embedded into paraffin. The paraffin sections (1 μm) were dewaxed, stained with hematoxylin and eosin, embedded in synthetic polymer (Dako Agilent, USA), and visualized using a DM6000B microscope (Leica, Germany) with 2.5-20x objectives. Two blinded independent experts assessed the violations of the structure of muscle fibers and signs of inflammation on microphotographs of longitudinal sections using a 3-point scale (no alterations, middle or strong alterations). The following parameters were examined: the diameter and shape of the muscle fibers; number and location of nuclei; any cellular infiltration (mostly by leucocytes); relative volume of interfiber space.

### RNA sequencing and data processing

A frozen muscle sample (∼15 mg) was lysed in 500 µl of an ExtractRNA buffer (Evrogen, Russia) using a drill homogenizer, and total RNA was extracted by a silica spin column (RNeasy Mini Kit, Qiagen, Germany). RNA concentration and integrity were evaluated by a fluorimetric assay (Qubit 4, ThermoScientific, USA) and capillary electrophoresis (TapeStation, Agilent, Germany), respectively. All samples had RIN >7. Strand-specific libraries were prepared by the NEBNext Ultra II Directional RNA Library Preparation kit (NEB, USA), as described previously (Makhnovskii et al., 2020), and sequenced (75 nucleotides, single end) with a median depth of 66 million reads per sample by NextSeq 550 (Illumina, USA). Raw data were deposited to NCBI GEO: GSE242202.

Sequencing quality was assessed using the FastQC tool (v.0.11.5). Low-quality reads and adapter sequences were deleted (Timmomatic tool, v0.36), and then the reads were aligned to the GRCh38.p13 primary genome assembly. Uniquely aligned reads were counted for known exons of each gene using the Rsubread package (R environment) and Ensembl annotation (GRCh38.101). Differential expression analysis was performed by the DESeq2 method (analysis of unpaired samples with the Benjamini-Hochberg correction). Differentially expressed genes (DEGs) were defined as protein-coding genes (also polymorphic pseudogene or translated pseudogene) with p_adj_ <0.01, |Fold Change| ≥1.25, and the expression level TPM >1 (Transcripts Per Million) was calculated using kallisto v0.46.2.

### Transcription factor binding site enrichment analysis (positional weight matrix [PWM] approach)

To predict transcription factors associated with primary aging genes, the individual promoter regions identified for human skeletal muscle were investigated by gene-specific positions of the transcription start site and open chromatin (Makhnovskii et al., 2022). Enrichment of the transcription factor binding sites (and corresponding transcription factors) in individual promoter regions of genes associated with primary aging was performed by the GeneXplain platform (the “Search for enriched TFBSs (tracks)” function http://wiki.biouml.org/index.php/Search_for_enriched_TFBSs_(tracks)_(analysis)) using the PWM database TRANSFAC v2022.2, as described elsewhere (Makhnovskii et al., 2022). Briefly, if a transcription factor has several PWM, the most enriched PWM was used. The maximum fold enrichment (FE_adj_, statistically corrected odds ratios with a confidence interval of 99%) was determined for each PWM (site frequency ≤1 per 2,000 bp) relative to that in 5,000 random individual promoters showing no differential expression in any of the experimental groups (DESeq2 method, p_adj_ >0.4). Adjusted fold enrichment (FE_adj_) >1 for both transcription factor binding site frequency and promoter’s sequence number (the binomial test and exact Fisher’s test, respectively), as well as FDR <0.05, were set as significance thresholds. The FE_adj_ evaluated by these tests demonstrated a strong correlation (Supplement Table S6). Only expressed transcription factors (TPM in OP >1) were included in further analysis.

### Quantitative shotgun proteomics and data processing

A frozen muscle sample (∼10 mg) was homogenized in 140 µl of a lysis buffer (5% SDS, 0.1 M DTT, 0.1 M TRIS, pH 7.6). Lysate was boiled (95 °C, 5 min), transferred to a microTUBE AFA vial, and sonicated (mean power 20 W, 30 s × 4) by a ME220 focused-ultrasonicator (Covaris, USA). After centrifugation (5 min, 30,000 g), protein concentration in the supernatant was measured by a fluorimetric assay (Qubit 4).

Protein hydrolysis was carried out using a S-Trap micro spin column (ProtiFi, USA). Samples (85 μg of proteins) diluted in the lysis buffer (final volume 24 μl) were alkylated (20 mM iodoacetamide, 15 min); further steps were performed according to the manufacturer’s protocol using proteases trypsin and Lys-C (2 h at 47° C, 1:15 and 1:30, respectively, Promega, USA). Peptides (20 μg) were dried down, resuspended in 100 mM triethylammonium bicarbonate, and labeled by isobaric tags for 1.5 h (TMT 10-plex or 16-plex kit, Thermo Scientific, USA). The reaction was quenched (0.3% hydroxylamine, 15 min), and the samples were combined.

The mixture of labeled peptides was subjected to reverse-phase LC fractionation using an Agilent 1200 Series HPLC (Agilent, USA) to obtain 24 fractions then fractions were combined (1 and 13, etc.) to obtain 12 fractions. Briefly, the peptides were concentrated on an Accalaim μ-Precolumn (0.5 mm × 3 mm, particle size 5 μm; Thermo Scientific) in the isocratic mode at a 10 μL/min flow for 5 min in the mobile phase C (2% acetonitrile, 0.1% formic acid). The peptides were then separated on a Peaky C18 column (100 μm × 300 mm, particle size 1.9 μm; Molecta, Russia) in a gradient mode of elution.

Each fraction was analyzed triple by an HPLC Ultimate 3000 RSLC nano system (Thermo Scientific) and a Q Exactive HF-X Hybrid Quadrupole-Orbitrap mass spectrometer (Thermo Scientific) using the nanoelectrospray ion source in the positive mode of ionization (Thermo Scientific). The gradient (90 min) was formed by the mobile phase A (0.1% formic acid) and B (80% acetonitrile, 0.1% formic acid) at a 0.4 μL/min flow. The ionizing voltage was 2.1 kV. MS spectra were acquired at a resolution of 60,000 in the 390–1300 m/z range; fragment ions were mass scanned at a resolution of 60,000 at the range from m/z 120 to the upper m/z value as assigned by the mass to charge state of the precursor ion. All tandem MS scans were performed on ions with a charge state from z = 2+ to z = 4+. Synchronous precursor selection facilitated the simultaneous isolation of up to 40 MS2 fragment ions. The maximum ion accumulation times were set to 50 ms for precursor ions and 25 ms for fragment ions. AGC targets were set to 10^6^ and 10^5^ for precursor ions and fragment ions, respectively.

The search and identification of peptides and proteins were carried out using the MaxQuant platform (2.1.4.0; Max Planck Institute of Biochemistry) at default settings (the FDR for peptides 1%, N-terminal acetylation and methionine oxidation as variable modifications and cysteine carbamidomethylation as fixed modification). Further analysis was performed using the Perseus platform (1.6.5; Max Planck Institute of Biochemistry). To avoid the batch effect related to labeling by the TMT 10-plex and 16-plex kits, the mean intensity of the reporter ions of all samples labeled by 10-plex isobaric tags was normalized to that of all samples labeled by 16-plex tags (Supplementary Figure S8). After filtration (removing potential contaminants, reverse peptides, peptides identified only by site), proteins identified by >1 unique+razor peptide and present in >70% of samples were taken for further analysis. To examine differentially expressed proteins, the one-way analysis of variance with q-value (Benjamini-Hochberg corrected p-value) <0.05 and post-hoc analysis (Tukey’s test; p<0.05) were applied.

### Statistical analysis

Physiological data are expressed as the median and interquartile range. The Kruskal-Wallis one-way analysis of variance with p-value <0.05 and post-hoc analysis (Dunn’s test; p<0.05) were used to compare groups.

The functional enrichment of biological processes and cellular components relative to a set of reference genes (all expressed [TPM >1] protein-coding genes) was performed for sets of up- and down-regulated genes or proteins by the DAVID 6.8 (p_adj_<0.05 Fisher exact test with Benjamini correction) using the UNIPROT KW BP/CC, KEGG PATHWAY, and GENE ONTOLOGY BP/CC DIRECT databases.

The protein-protein interaction network (physical subnetwork) was generated by the String database 12.0 with interaction sources Textmining, Experiments, and Databases using the MCL clustering method.

To study the intergroup expression trajectory (YH-YP-OP) of differentially expressed proteins belonging to enriched functional categories and corresponding mRNAs, the normalized expression (*z*-score) of each protein/mRNA was used.

We compared the primary ageing genes identified in our study with genes that changed expression in *i*) skeletal muscle and other tissues in older individuals/animals (meta-analyses of transcriptomic data (Avelar et al., 2020; Palmer et al., 2021; Su et al., 2015) and *ii*) senescent human fibroblasts (Casella et al., 2019), as well as with *iii*) genes previously associated with aging and/or senescence in muscle or other tissues (a Medline search). For the Medline search we used terms: ‘(gene name) AND (aging OR senescence) AND muscle’ and ‘(gene name) AND (aging OR senescence)’. Articles showing a mechanistic link between gene expression and the aging or senescence phenotype in skeletal/cardiac muscle tissue/muscle cells or other tissues/cells were selected. Studies that showed an association with age-related diseases or cancer were declined.

## Acknowledgments

The authors acknowledge the clinicians and laboratory staff at Medical Research and Educational Center of Lomonosov Moscow State University for their excellent assistance in human biomaterial collection and are grateful for the opportunity to use the mass spectrometry equipment of the “Human Proteome” Core Facility (Institute of Biomedical Chemistry, Moscow).

## Authors’ contributions

Conceptualization, A.Yu. E. and D.V.P.; methodology, V.G.Z., A.Yu. E., and D.V.P.; formal analysis, N.S.K., M.A.O., M.A.V., T.F.V., V.G.Z., N.E.V., P.A.M., O.A.G., Y.R.B., V.V.P., E.M.L., F.A.K., A.Yu. E., and D.V.P.; investigation, N.S.K., M.A.O., M.A.V., T.F.V., V.G.Z., N.E.V., P.A.M., O.A.G., Y.R.B., V.V.P., E.M.L., F.A.K., A.Yu. E., and D.V.P.; resources, P.A.M. and D.V.P.; data curation, P.A.M. and D.V.P.; writing—original draft preparation, N.S.K. and D.V.P.; writing—review and editing, N.S.K., M.A.O., P.A.M., A.Yu. E. and D.V.P.; project administration, A.Yu. E. and D.V.P. All authors have read and agreed to the published version of the manuscript.

## Data availability statement

RNA-seq datasets generated for this study can be found in NCBI Gene Expression Omnibus (GEO) under accession number GSE242202; all processed proteomic data can be found in supplementary files.

## Supporting Information

**Supplementary Figure S1. Pathology- and age-related physiological changes in young (YP) and older (OP) patients compared to young healthy (YH) individuals.**

The red dotted lines indicate the normal range. YH: n = 15, YP: n = 8, OP: n = 37. * – difference from YH and # – difference from YP at p <0.001.

**Supplementary Figure S2. Overlapping of sets of differentially expressed genes.**

Proteins (as well mRNAs) that unidirectionally change expression in both OP *vs.* YP and OP *vs.* YH were defined as genes associated with primary aging – shaded overlapping area (then mRNAs whose protein products did not change expression were excluded: see also Figure 4 and Supplementary Table S5).

**Supplementary Figure S3. Intergroup expression trajectory (YH-YP-OP) of differentially expressed mRNAs related to inflammatory and immune response.**

Normalized expression (*z*-score) for each gene, as well as the name and ID of functional categories are indicated. YH: n = 15, YP: n = 8, OP: n = 37. *** and ### – difference at p <0.001 from YH and YP, respectively.

**Supplementary Figure S4. Comparison of the primary aging genes identified in our study with literature data.**

A – The set of primary aging genes was significantly enriched (Fisher’s exact test) in genes that changed expression in skeletal muscle of older individuals/animals, but not in other tissues (meta-analyses of transcriptomic data (Avelar et al., 2020; Palmer et al., 2021; Su et al., 2015); overlapped 21% and 13%, respectively). OR – odds ratio.

B – The set of primary aging genes showed no significant enrichment (Fisher’s exact test) in genes induced by replicative aging in human fibroblasts (Casella et al., 2019). OR – odds ratio. C – A Medline search indicated that the set of primary aging genes overlapped with genes previously associated with aging and/or senescence in muscle or other tissues (13% and 25%, respectively).

**Supplementary Figure S5. Sex-related differences in gene expression in the older patient group.**

A and C – Sex-related genes/proteins in the OP group showed little/no overlapping with OP *vs.* YH and OP *vs.* YP.

B and D – The principal component analysis showed that male and female transcriptomic (B) and proteomic (D) profiles were quite similar. Empty circles/asterisks indicate females.

**Supplementary Figure S6. Dependence of statistical power on sample size (SS) in older and young patients and on gene expression (normalized read count).**

A – Comparison OP *vs.* YH (n = 3, 7, 15, and 37 for OP (data simulation) and n = 15 for YH).

B – Comparison YP *vs.* YH (n = 3, 7, 15, and 37 for YP (data simulation) and n = 15 for

**Supplementary Figure S7. Dependence of the number of differentially expressed genes on sample size and on various cutoff criteria (OP *vs.* YH).**

n = 15 for YH, n = 3 to 37 for OP.

**Supplementary Figure S8. Additional normalization of reporter ion intensity.**

To avoid the batch effect related to labeling by the TMT 10-plex and 16-plex kits (top), the mean intensity of the reporter ions of all samples labeled by 10-plex isobaric tags was normalized to that of all samples labeled by 16-plex tags (bottom).

**Supplementary Table S1. All expressed (TPM >1) and differentially expressed (p_adj_ <0.01, |Fold Change| ≥1.25) mRNAs.**

**Supplementary Table S2. Function enrichment analysis of differentially expressed mRNAs (p_adj_ <0.01, |Fold Change| ≥1.25) relative to all expressed mRNAs (TPM >1).**

**Supplementary Table S3. All detected and differentially expressed proteins (p_adj_ <0.05) and their reporter ion intensity.**

**Supplementary Table S4. Function enrichment analysis of differentially expressed proteins (p_adj_ <0.05) relative to all detected proteins. Sets of proteins with different patterns of regulation at the mRNA level were analyzed separately (see also Figure 3)**.

**Supplementary Table S5. Genes associated with primary aging and their functions.**

**Supplementary Table S6. Transcriptional factors associated with primary aging genes (all results of the transcription factor binding site enrichment analysis).**

For transcription factor prediction, adjusted fold enrichment (FE_adj_) >1 for both transcription factor binding site frequency and promoter’s sequence number (the binomial test and exact Fisher’s test, respectively), as well as FDR <0.05, were set as significance thresholds.

