## Supplementary Figure for "Age-related changes in human skeletal muscle transcriptome and proteome are more affected by chronic inflammation and physical inactivity than primary aging"

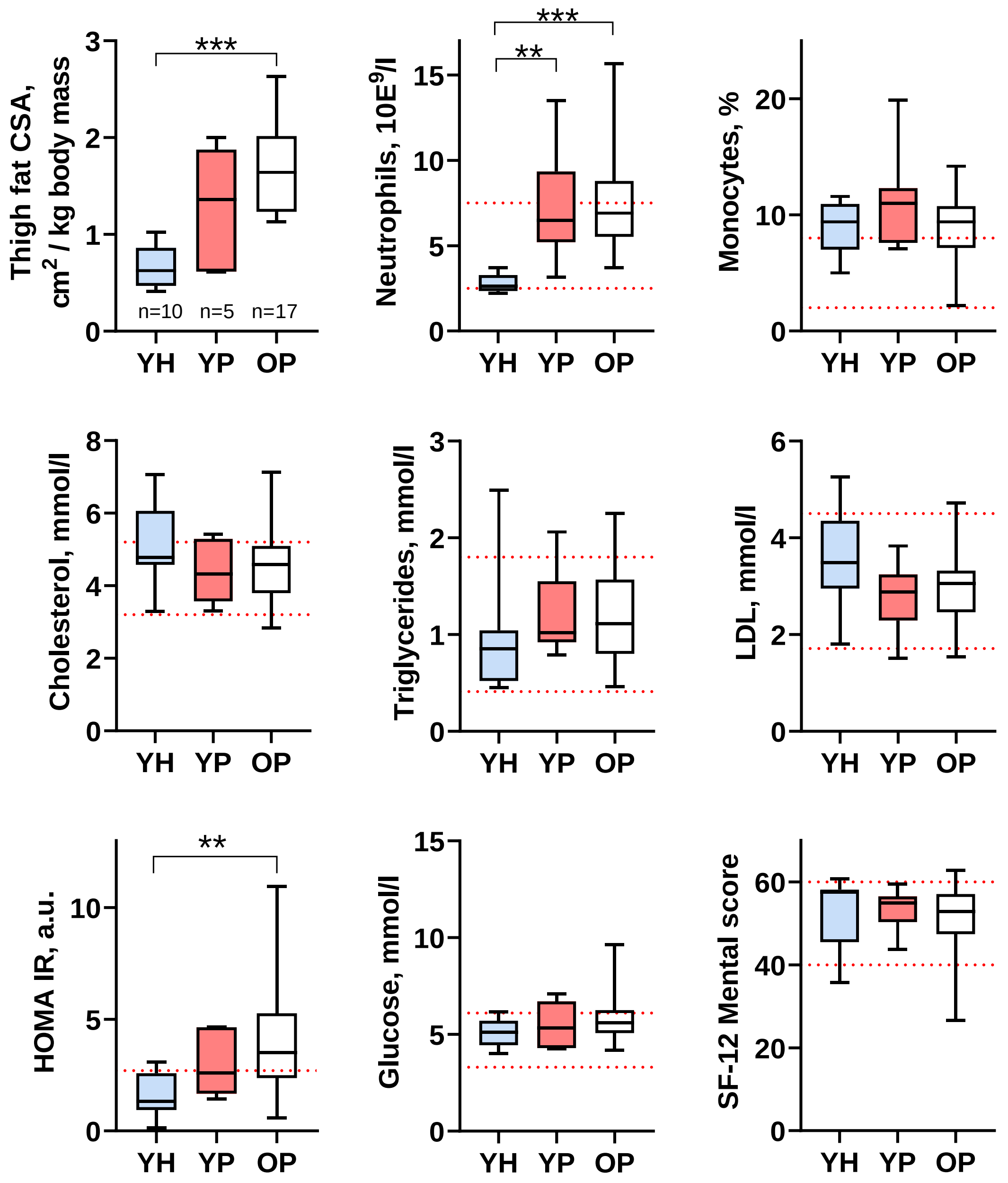


**Supplementary Figure S1. Pathology- and age-related physiological changes in young (YP) and older (OP) patients compared to young healthy (YH) individuals.**

The red dotted lines indicate the normal range. YH: n = 15, YP: n = 8, OP: n = 37. * – difference from YH and # – difference from YP at p <0.001.

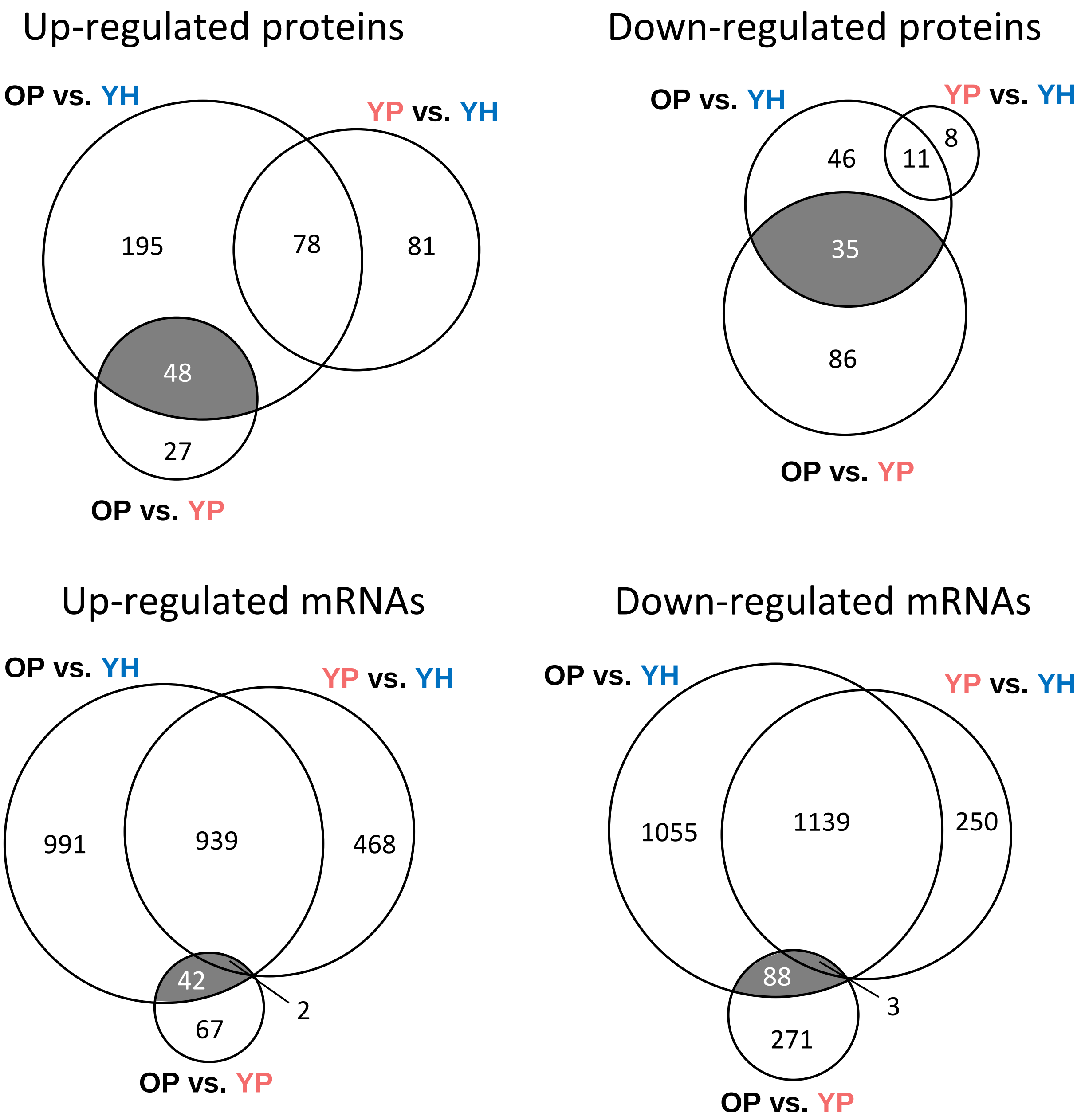


**Supplementary Figure S2. Overlapping of sets of differentially expressed genes.**

Proteins (as well mRNAs) that unidirectionally change expression in both OP *vs.* YP and OP *vs.* YH were defined as genes associated with primary aging – shaded overlapping area (then mRNAs whose protein products did not change expression were excluded: see also Figure 4 and Supplementary Table S5).

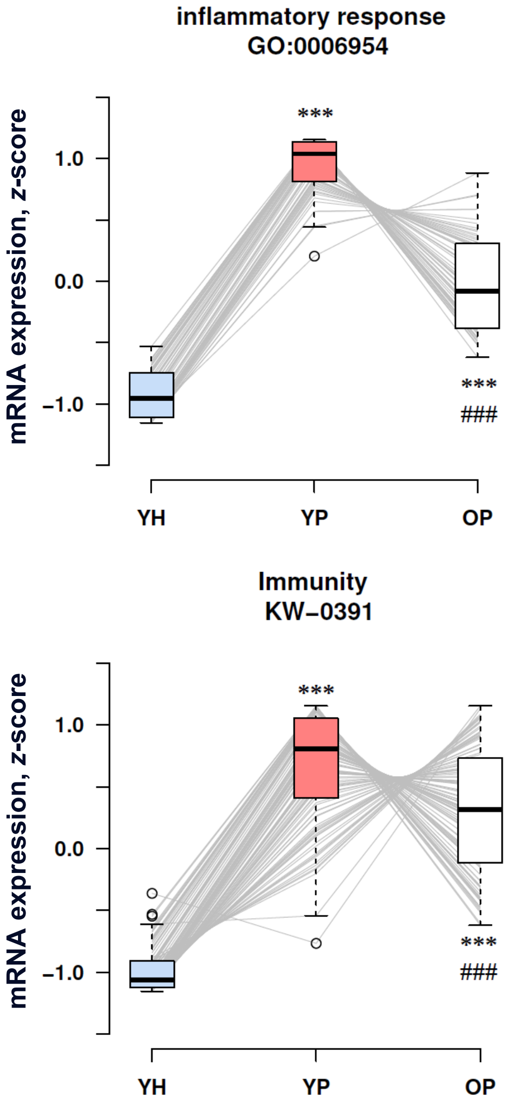


**Supplementary Figure S3. Intergroup expression trajectory (YH-YP-OP) of differentially expressed mRNAs related to inflammatory and immune response.**

Normalized expression (*z*-score) for each gene, as well as the name and ID of functional categories are indicated. YH: n = 15, YP: n = 8, OP: n = 37. *** and ### – difference at $p < 0.001$ from YH and YP, respectively.

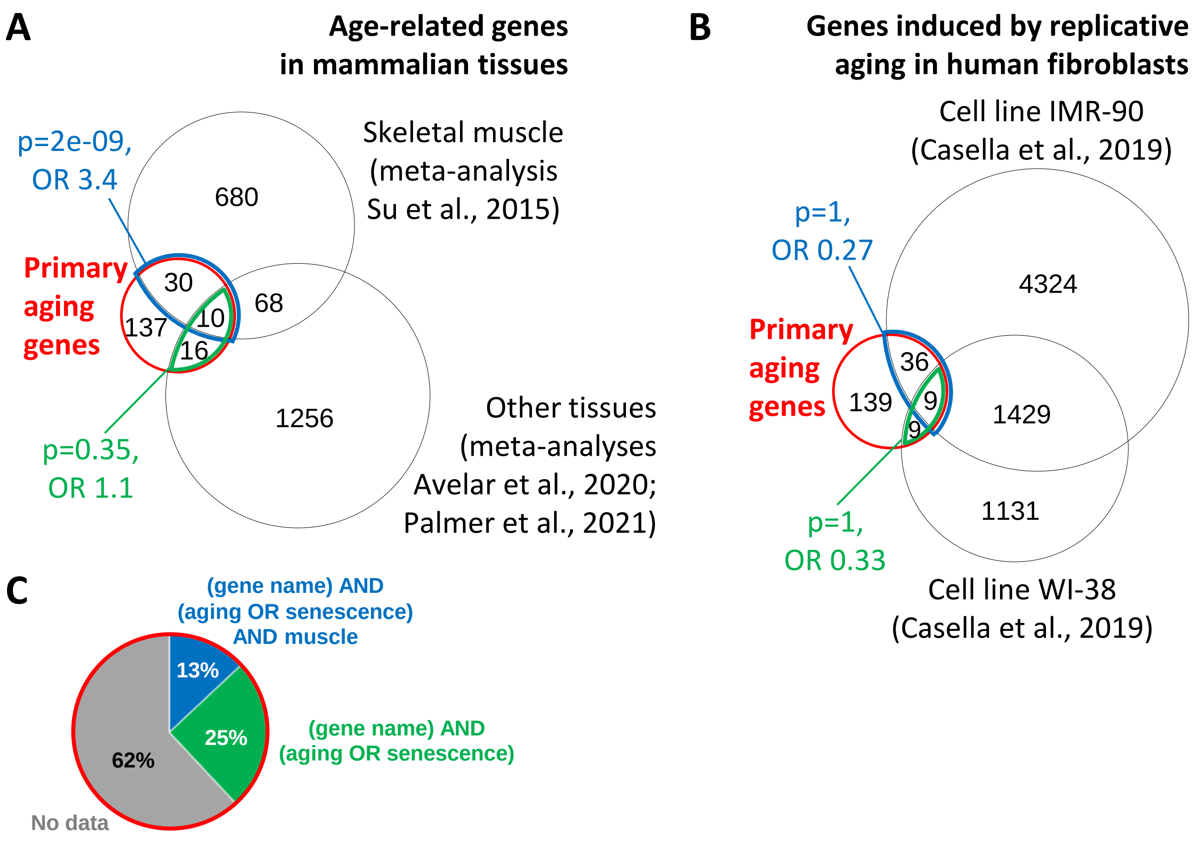


**Supplementary Figure S4. Comparison of the primary aging genes identified in our study with literature data.**

A – The set of primary aging genes was significantly enriched (Fisher's exact test) in genes that changed expression in skeletal muscle of older individuals/animals, but not in other tissues (meta-analyses of transcriptomic data (Avelar et al., 2020; Palmer et al., 2021; Su et al., 2015); overlapped 21% and 13%, respectively). OR – odds ratio.

B – The set of primary aging genes showed no significant enrichment (Fisher's exact test) in genes induced by replicative aging in human fibroblasts (Casella et al., 2019). OR – odds ratio.

C – A Medline search indicated that the set of primary aging genes overlapped with genes previously associated with aging and/or senescence in muscle or other tissues (13% and 25%, respectively).

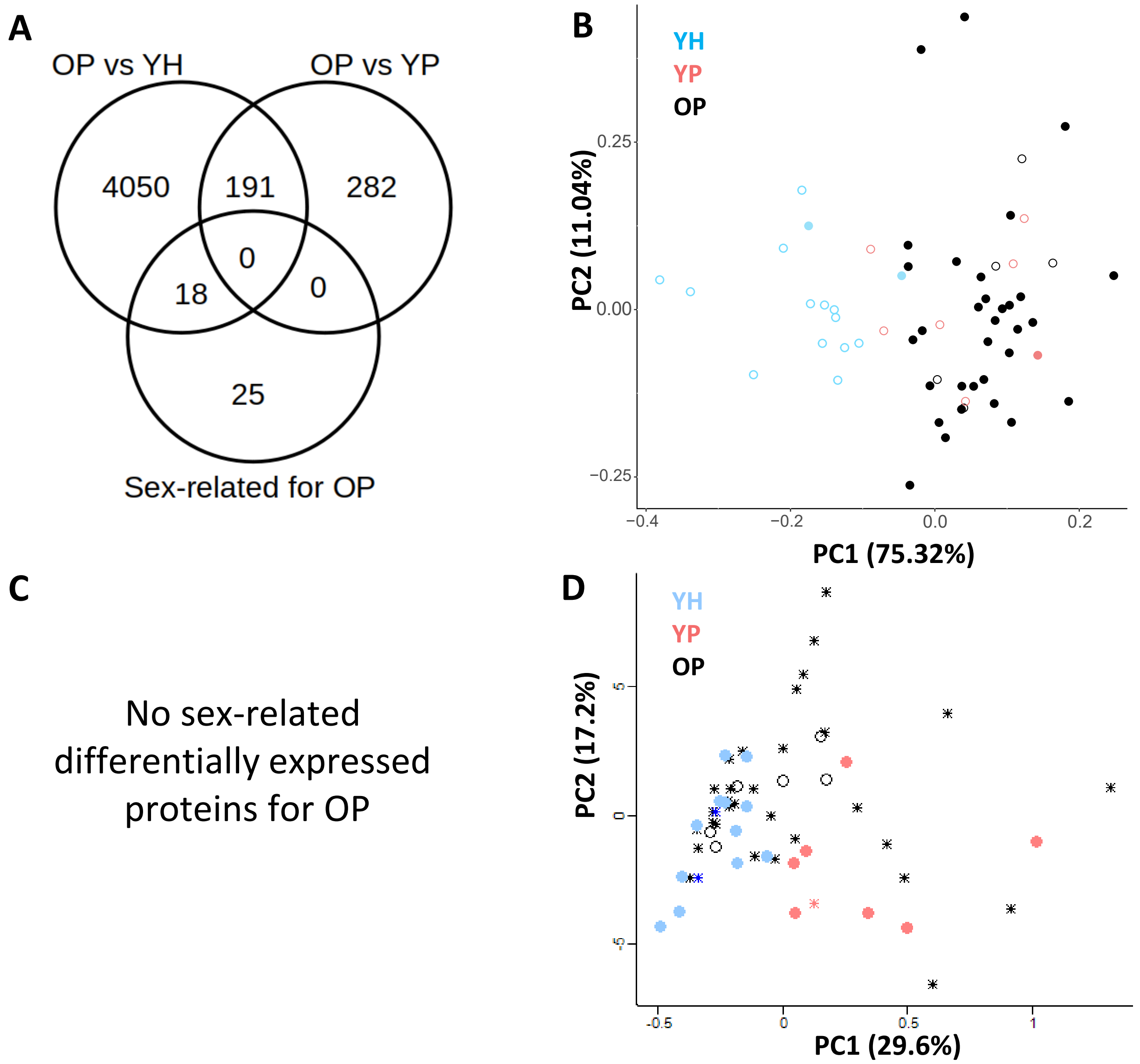


**Supplementary Figure S5. Sex-related differences in gene expression in the older patient group.**

A and C – Sex-related genes/proteins in the OP group showed little/no overlapping with OP vs. YH and OP vs. YP.

B and D – The principal component analysis showed that male and female transcriptomic (B) and proteomic (D) profiles were quite similar. Empty circles/asterisks indicate females.

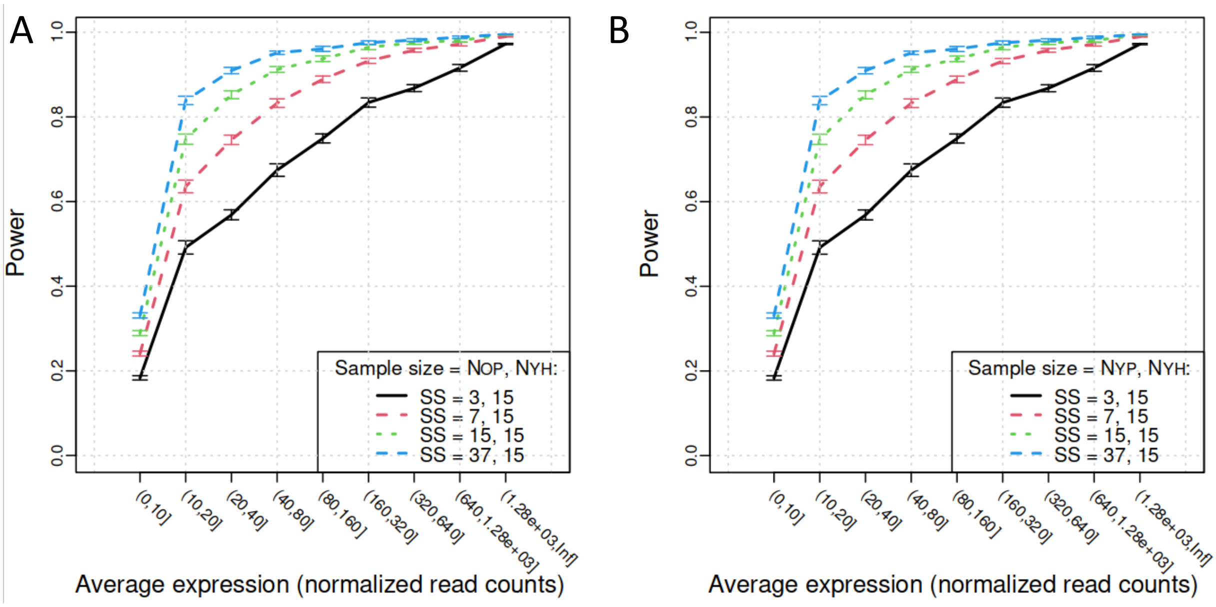


**Supplementary Figure S6. Dependence of statistical power on sample size (SS) in older and young patients and on gene expression (normalized read count).**

A – Comparison OP vs. YH (n = 3, 7, 15, and 37 for OP (data simulation) and n = 15 for YH).

B – Comparison YP vs. YH (n = 3, 7, 15, and 37 for YP (data simulation) and n = 15 for YH).

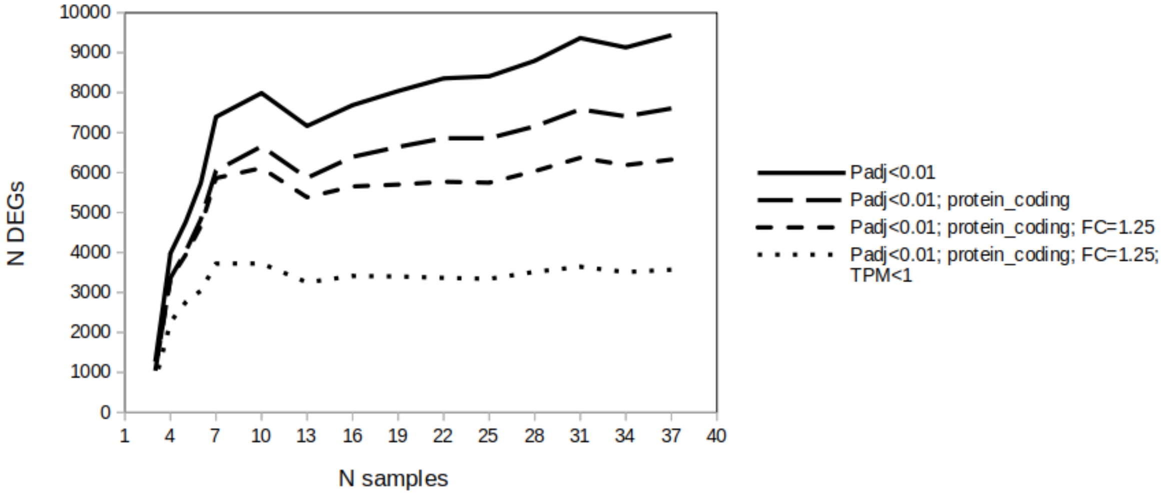


**Supplementary Figure S7. Dependence of the number of differentially expressed genes on sample size and on various cutoff criteria (OP vs. YH).**

n = 15 for YH, n = 3 to 37 for OP.

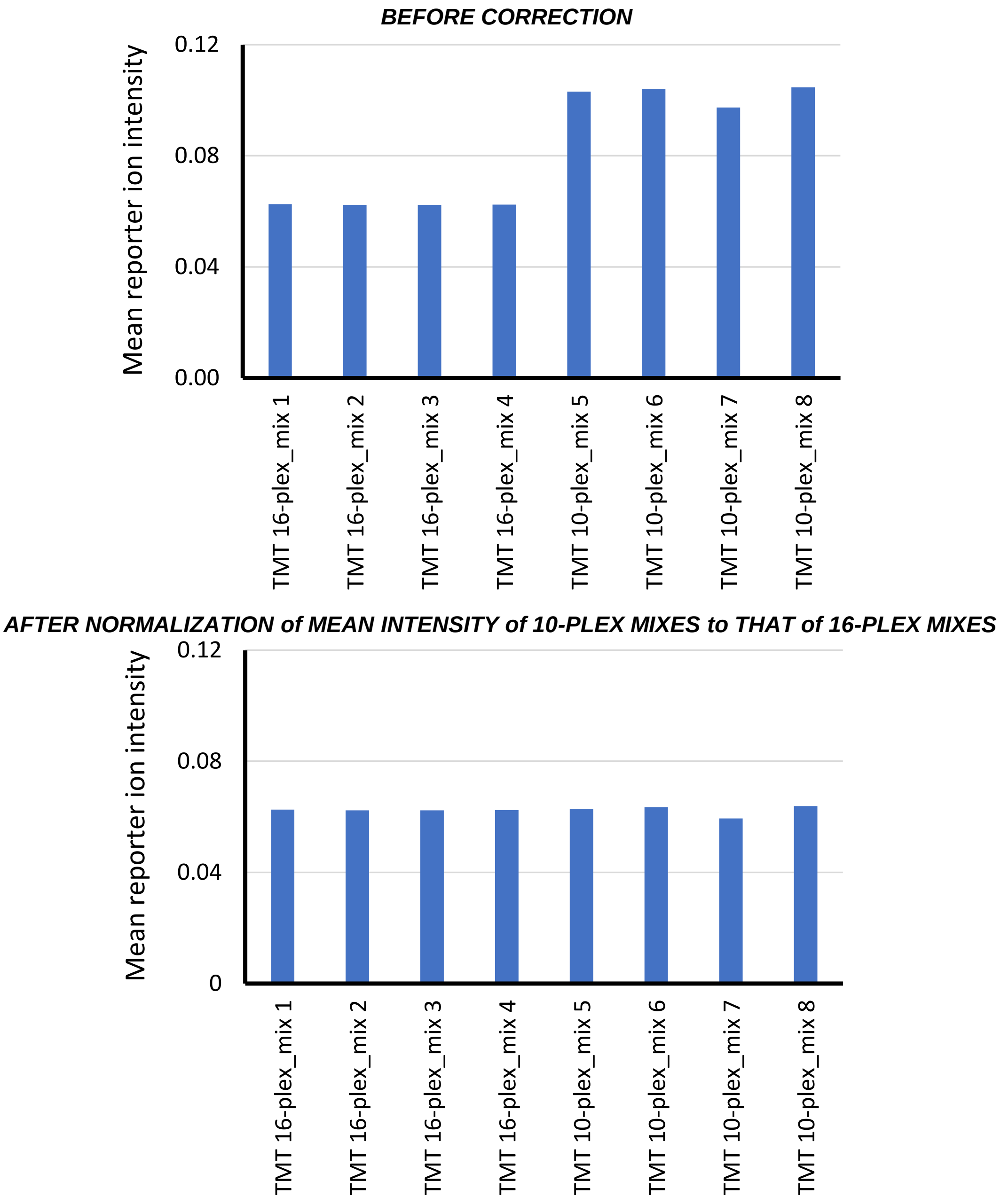


**Supplementary Figure S8. Additional normalization of reporter ion intensity.**

To avoid the batch effect related to labeling by the TMT 10-plex and 16-plex kits (top), the mean intensity of the reporter ions of all samples labeled by 10-plex isobaric tags was normalized to that of all samples labeled by 16-plex tags (bottom).
